# Effective structure-aware protein alignment via residue-level contrastive learning

**DOI:** 10.1101/2024.03.09.583681

**Authors:** Ronghui You, Ziye Wang, Kunpeng Liu, Wei Zheng, Qiqige Wuyun, Yuhao Yi, Shanfeng Zhu

## Abstract

Protein alignment is indispensable for biological discovery, supporting structure comparison, functional annotation, and evolutionary inference. While structure-based methods are highly effective at detecting structural similarity, their applicability is constrained by the limited availability of experimentally resolved protein structures and high computational cost. Sequence-based approaches using pretrained protein language models (pLMs) provide scalable alternatives, yet supervised methods based on differentiable dynamic programming have not consistently outperformed simpler unsupervised strategies. Here, we present CLAlign, a structure-aware protein alignment framework based on contrastive learning. CLAlign fine-tunes a pretrained pLM to generate structure-aware residue-level embeddings enriched with structural context, without relying on differentiable dynamic programming. It represents the first supervised approach to consistently outperform unsupervised pLM-based methods, and it naturally extends to both sequence- and structure-based alignment by flexibly adopting different protein language model encoders. CLAlign achieves state-of-the-art accuracy on the MALIDUP and MALISAM benchmarks, outperforming existing sequence-based methods by large margins while remaining highly efficient. Moreover, its alignment scores show clear biological interpretability: in remote homology detection on SCOPe, CLAlign performs comparably to structure-based methods such as TM-align while far exceeding all sequence-based baselines. Together, these results establish CLAlign as a simple, extensible, and biologically meaningful framework for protein alignment.

## 1 Introduction

Protein alignment is a foundational task in bioinformatics, underpinning sequence comparison, structure analysis, functional annotation, and evolutionary inference [1]. By identifying conserved residue correspondences, alignment supports the transfer of functional knowledge to uncharacterized proteins, the interpretation of disease-associated variation, and the prioritization of sites for mutational studies or drug design [2]. At larger scales, alignment enables evolutionary analyses by tracing the lineage, duplication, and adaptation of protein families across species and deep evolutionary time [2].

Protein alignment methods are commonly grouped by input type into structure-based and sequence-based approaches. Structure-based methods generally provide highly accurate residue mappings by explicitly optimizing geometric consistency. Representative methods include FAST, which accelerates structural comparison using geometric hashing [3]; TM-align, which identifies optimal superpositions by maximizing the TM-score [4]; Dali, which detects conserved inter-residue distance patterns [5]; and GTAlign, which enables fast search and alignment through spatial indexing [6]. Despite their effectiveness in detecting structural similarity, structure-based methods remain constrained by both data availability and scalability. Most proteins lack experimentally determined structures, and even high-quality protein structure prediction methods [7–10] can be unreliable for short, low-homology, or otherwise challenging targets. In addition, many structure-based workflows rely on computationally iterative optimization and rigid-body superposition, which limits throughput for large-scale searches. Sequence-based alignment is therefore indispensable for proteome-scale studies, but it must infer structural correspondence from sequence alone, particularly at low sequence identity, where classical substitution matrices and hand-crafted scoring schemes lose discriminative power. Bridging this gap, retaining the scalability of sequence-based input while improving structure faithfulness, remains a central challenge in protein alignment.

Recently, protein language models (pLMs) [11–15], exemplified by ESM, have advanced rapidly and demonstrated strong performance across a wide range of downstream bioinformatics tasks. Associated with deep learning, pLMs have revitalized sequence-based alignment by providing residue embeddings that capture contextual and, to some extent, structural signals [16–20]. These pLM-based sequence alignment methods can be broadly divided into unsupervised and supervised approaches. Unsupervised methods such as pLM-BLAST [16], PLMAlign [17], and EBA [18] compute residue-residue similarity directly from pretrained embeddings (e.g., using cosine or dot-product similarity) and then apply classical dynamic programming (DP), including Smith-Waterman (SW) [21] or Needleman-Wunsch (NW) [22], to recover alignments. Despite their simplicity, these approaches often perform strongly, consistent with the notion that large pLMs encode structural information even without task-specific fine-tuning. However, because these embeddings are not optimized for alignment, similarity matrices can be noisy and difficult for dynamic programming to interpret, particularly for analogous structures with weak sequence similarity. Supervised alternatives, including DeepBLAST [19] and DEDAL [20], instead seek to learn structure-aware embeddings through differentiable dynamic programming. In practice, differentiable DP increases implementation and optimization complexity and can be unstable during training. Moreover, reported evaluations show that these supervised DP-based frameworks do not consistently surpass strong unsupervised pLM baselines [17, 19, 20]. Together, these observations motivate training objectives that directly shape the embedding space to make true residue correspondences more separable, while preserving the simplicity and efficiency of standard dynamic programming at inference time.

This study presents CLAlign, a structure-aware protein alignment framework built on contrastive learning. CLAlign fine-tunes a pretrained pLM to produce residue-level representations enriched with structural context and then uses these representations to derive a substitution matrix that can be directly consumed by standard dynamic programming. In this design, learning occurs entirely in the embedding space, where contrastive supervision increases similarity for aligned residue pairs and decreases similarity for non-corresponding positions, thereby sharpening the alignment signal used by downstream dynamic programming. The framework is designed to accommodate either sequence or structure encoders, enabling a unified treatment of sequence alignment and structure alignment within a single objective. Across benchmarks, CLAlign achieves state-of-the-art accuracy among sequence-based methods and, to our knowledge, provides the first supervised approach that consistently outperforms unsupervised pLM-based aligners such as pLM-BLAST and PLMAlign. In addition to improved alignment quality, CLAlign yields alignment scores that track evolutionary homology more faithfully, supporting biologically interpretable measures of relatedness that are useful for search, ranking, and downstream comparative analyses.

## 2 Results

### 2.1 Pairwise protein sequence alignment with CLAlign

We introduce CLAlign, a unified protein alignment framework based on contrastive learning (Fig. 1). CLAlign fine-tunes a pretrained protein language model (pLM) to learn structure-aware residue representations by bringing embeddings of aligned residues closer while pushing those of non-aligned residues apart. The resulting embeddings are used to construct a substitution matrix via cosine similarity, which is then consumed by standard dynamic programming algorithms, including Needleman–Wunsch for global alignment and Smith–Waterman for local alignment.

**Figure 1.**
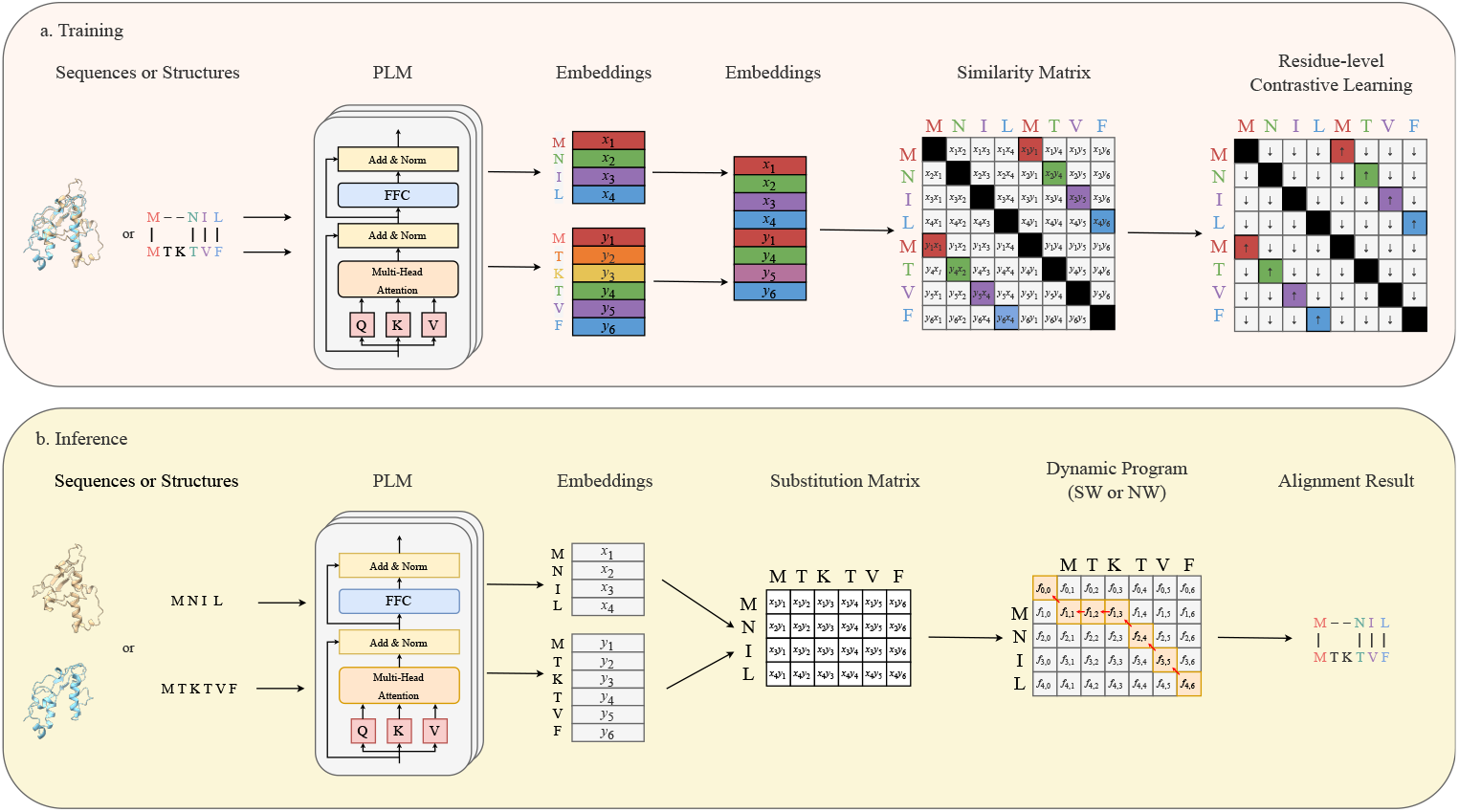
Overview of CLAlign. The upper panel illustrates the training stage: a pair of aligned protein sequences or structures is encoded by a pLM to generate residue embeddings. Cosine similarities are computed between every pair of residues in the aligned regions (including pairs within the same sequence), with diagonal entries ignored, forming the similarity matrix. Contrastive learning fine-tunes the pLM to refine the embeddings by pulling matched residue pairs closer and pushing non-matching pairs apart, which corresponds to increasing similarity values for colored cells and decreasing those for white cells in the similarity matrix. Cells on the diagonal (shown in black) are ignored and not involved in the optimization. The lower panel illustrates the inference stage: given two sequences or structures, a substitution matrix is obtained using the same process as in training with the fine-tuned pLM. The substitution matrix is then processed by the Smith–Waterman or Needleman–Wunsch algorithm to produce the final alignment.

In contrast to supervised approaches based on differentiable dynamic programming, CLAlign learns structural correspondence directly in embedding space, yielding a simpler and more scalable training objective. The framework is also extensible since alternative encoders can be substituted without changing the contrastive objective, enabling both sequence-only and structure-informed alignment within the same architecture. Detailed information could be found in the Methods Section.

### 2.2 CLAlign improves alignment accuracy

Alignment accuracy was assessed on two widely used datasets, MALIDUP [23] and MALISAM [24]. MALIDUP comprises 241 homologous domain pairs derived from internal duplications, whereas MALISAM contains 130 structurally analogous motifs lacking common ancestry. Both datasets provide expert-curated reference alignments, in which aligned residue pairs are defined using structural, evolutionary and functional considerations rather than sequence identity alone. CLAlign was compared against seven sequence-based alignment methods spanning three categories: (i) classical dynamic programming methods (Needleman-Wunsch and Smith-Waterman), (ii) unsupervised pLM-based methods (pLM-BLAST [16], PLMAlign [17], and EBA [18]), and (iii) supervised pLM-based methods (DEDAL [20] and DeepBLAST [19]).

PLM-BLAST, PLMAlign, EBA, and DeepBLAST use ProtT5 [14] as their encoder, whereas DEDAL relies on its own pretrained model. For CLAlign, we evaluated two encoder variants, ProtT5 and ProstT5 [15], the latter incorporating structural information during pretraining to produce more structure-aware representations. Alignment quality was quantified primarily by F1-score, which summarizes the trade-off between precision and recall at the level of individual residue pairs. Precision and recall are also reported to facilitate interpretation of whether gains arise from fewer spurious matches, increased coverage of true matches, or both.

Table 1 shows that CLAlign achieves the highest F1-scores on both datasets. With ProtT5, CLAlign improves upon the strongest baseline, PLMAlign, by 15.2% on MALIDUP and 45.1% on MALISAM, indicating that contrastive fine-tuning enhances structure awareness in pLM embeddings. The larger gain on MALISAM, which involves structurally analogous but evolutionarily unrelated proteins, suggests that CLAlign learns correspondence that goes beyond sequence similarity, a central limitation of unsupervised approaches. Substituting ProtT5 with the more structure-aware ProstT5 further increases performance, yielding additional gains of 4.8% on MALIDUP and 12.0% on MALISAM. These results demonstrate that the contrastive-learning objective is compatible with, and benefits from, stronger pretrained encoders.

**Table 1.** Performance comparison of protein alignment on MALIDUP and MALISAM.

| Methods | encoder | MALIDUP |  |  | MALISAM |  |  |
| --- | --- | --- | --- | --- | --- | --- | --- |
|  |  | Precision | Recall | F1-score | Precision | Recall | F1-score |
| NW | - | 0.197 | 0.121 | 0.150 | 0.043 | 0.026 | 0.032 |
| SW | - | 0.197 | 0.124 | 0.151 | 0.044 | 0.027 | 0.033 |
| pLM-BLAST | ProtT5 | 0.612 | 0.500 | 0.548 | 0.203 | 0.152 | 0.173 |
| PLMAlign | ProtT5 | 0.582 | 0.542 | 0.561 | 0.193 | 0.177 | 0.184 |
| EBA | ProtT5 | 0.548 | 0.525 | 0.536 | 0.160 | 0.137 | 0.147 |
| DeepBLAST | ProtT5 | 0.529 | 0.509 | 0.518 | 0.157 | 0.148 | 0.152 |
| DEDAL | DEDAL | 0.495 | 0.384 | 0.417 | 0.095 | 0.062 | 0.070 |
| CLAlign | ProtT5 | <u>0.642</u> | <u>0.650</u> | <u>0.646</u> | <u>0.261</u> | <u>0.274</u> | <u>0.267</u> |
| CLAlign | ProstT5 | **0.674** | **0.680** | **0.677** | **0.294** | **0.306** | **0.299** |

Supervised differentiable-DP methods (DeepBLAST and DEDAL) underperform the unsupervised pLM-based approaches on both datasets. By contrast, CLAlign enforces pairwise matching consistency through contrastive supervision, directly shaping the embedding space so that true correspondences are easier to recover by standard dynamic programming. This distinction helps explain an empirical gap in prior work, in which supervision alone did not consistently translate into improvements over strong unsupervised pLM baselines.

### 2.3 CLAlign improves structural consistency

Although F1-score serves as the primary evaluation metric for residue-level accuracy, it does not fully capture the structural plausibility of alignments. To further assess how well the alignments preserve protein structural organization, we complemented F1-score with two additional perspectives: TM-score and HEC (Helix–Sheet–Coil)-based secondary-structure consistency metrics, which together provide complementary perspectives on global and local structure preservation.

Figure 2a-b compares TM-scores obtained from sequence-based alignments by pLM-BLAST, PLMAlign, EBA, DeepBLAST and CLAlign (CLAlign-ProtT5 for a fair comparison). Each point corresponds to a protein pair; the x-axis shows the TM-score achieved by the manual reference alignment, and the y-axis shows the TM-score resulting from the compared method. Across both MALIDUP and MALISAM datasets, CLAlign attains the highest average TM-scores (0.496 and 0.280, respectively), exceeding the second-best method by 13.2% on MALIDUP and 22.3% on MALISAM. For most protein pairs, CLAlign produces higher TM-scores than competing methods, indicating that the structure-aware embeddings learned through contrastive fine-tuning produce alignments that better reflect the global structural similarity of proteins. These results indicate that the contrastive learning framework not only improves residue-level correspondences but also aligns the embedding space with true structural relationships at the fold level.

**Figure 2.**
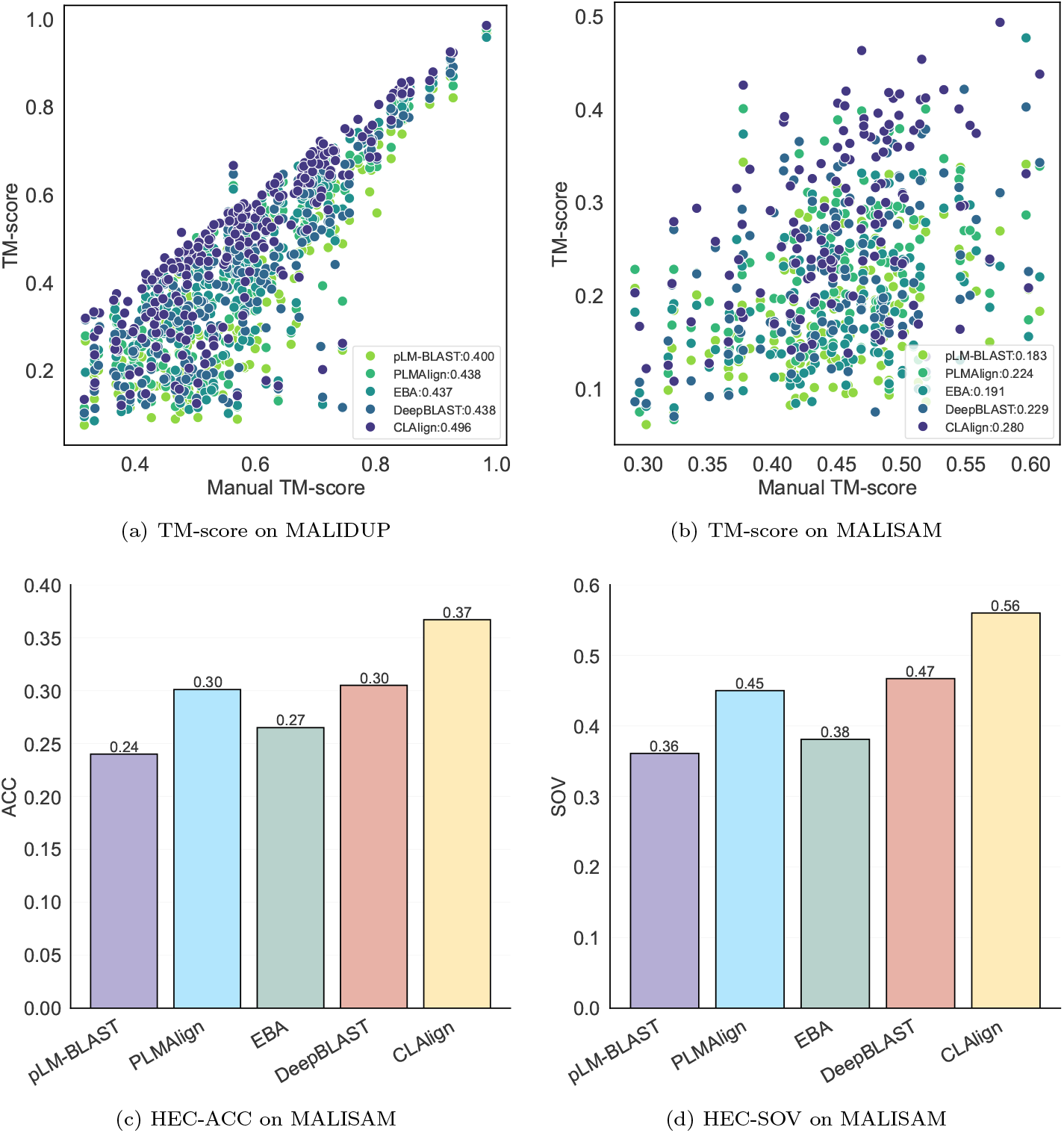
Comparison of alignment performance among different sequence-based methods on the MALIDUP and MALISAM datasets. Scatter plots (a) and (b) show the global structural similarity measured by TM-score, where each point represents one protein pair: the x-axis is the TM-score of the manual alignment, and the y-axis is the TM-score of the compared method. Bar plots (c) and (d) show the average secondary structure accuracy (ACC) and segment overlap score (SOV) on MALISAM, respectively, for each method.

Local secondary-structure preservation was assessed using two HEC-based measures: HEC-ACC, the per-residue agreement of secondary-structure states, and HEC-SOV, a segment overlap score. As shown in Fig. 2c-d, CLAlign achieves the strongest secondary-structure consistency on MALISAM (HEC-ACC = 0.367 and HEC-SOV = 0.560), corresponding to relative improvements of up to 12.7% and 19.9%, respectively. These gains indicate more faithful preservation of local folding patterns and secondarystructure continuity, particularly in the low-sequence-similarity regime represented by MALISAM. Because HEC evaluates the coherence of aligned local elements rather than overall geometry, improvements in HEC-ACC and HEC-SOV directly reflect improved matching of structurally analogous regions.

Overall, TM-score and HEC-based analyses indicate that contrastive learning in CLAlign improves structure awareness across multiple scales, enhancing global fold consistency while simultaneously strengthening the local coherence of aligned secondary-structure elements, information that is only partially captured by TM-score alone.

### 2.4 CLAlign improves distant homology detection

An alignment score is most useful when it not only ranks alignments correctly but also reflects biological relatedness. To test whether CLAlign’s scores capture evolutionary homology, distant homology detection was evaluated on the SCOPe40-test benchmark from PLMAlign [17]. SCOPe40-test contains 2,207 proteins. During the benchmark, each protein was used as a query against all targets, including itself, resulting in all-versus-all scores. Pairs assigned to different SCOPe folds were treated as negatives, following the benchmark protocol. CLAlign was compared with three sequence-based methods (pLM-BLAST, PLMAlign, and EBA) and with two structure-based alignment methods (TM-align and GTAlign). Performance was assessed following the SCOPe search-sensitivity protocol used by Foldseek and PLMSearch [25, 17]. For the ROC-based metric, targets for each query were ranked by alignment score, and sensitivity was measured by the area under the cumulative ROC curve up to the first false positive, following Foldseek. For all three ROC-based evaluations, false positives were consistently defined as targets from different SCOPe folds. To avoid confusion with conventional full-range AUROC, we refer to this metric as AUC_1FP_. At the family, superfamily, and fold levels, true positives were defined as proteins from the same family, from the same superfamily but not the same family, and from the same fold but not the same superfamily, respectively. MAP and P@k were computed using the same-fold proteins as positives.

As shown in Fig. 3, CLAlign achieves the best performance among all sequence-based approaches. It improves over the next-best sequence method by 18.3% and 52.3% in AUC_1FP_ at the superfamily and fold levels, respectively, and by 10.5% in MAP, while approaching the performance of structure-based alignment, TM-align, and GTAlign.

**Figure 3.**
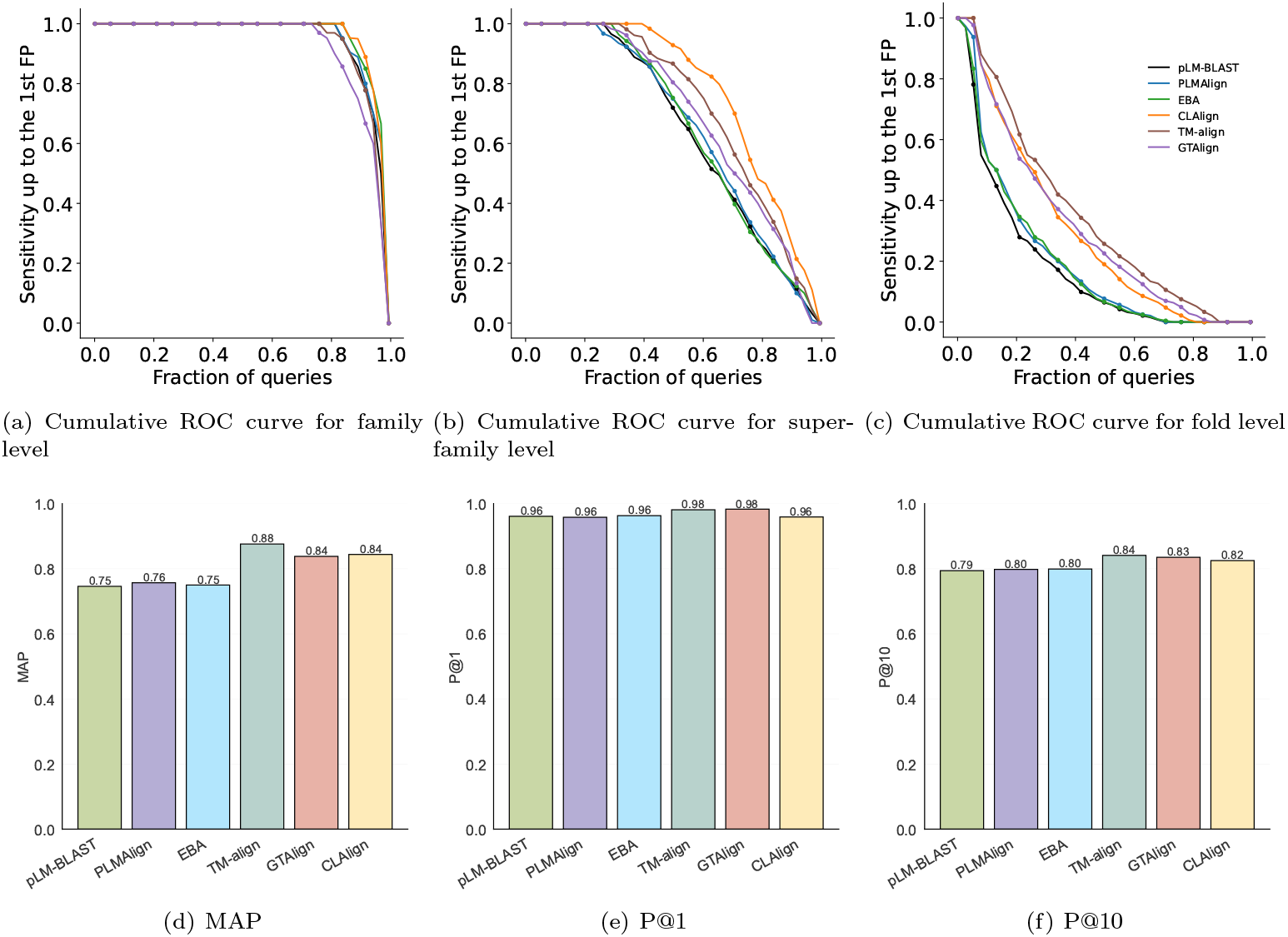
Cumulative ROC curves and summary metrics comparing the performance of pLM-BLAST, PLMAlign, EBA, CLAlign, TM-align, and GTAlign on the SCOPe40-test dataset. Panels (a–c) show cumulative ROC curves evaluated up to the first false positive at the family, superfamily, and fold levels, respectively. The corresponding areas are reported as AUC_1FP_. Panels (d–f) show bar plots for MAP, P@1, and P@10.

These results indicate that CLAlign’s alignment scores reliably distinguish homologous from nonhomologous pairs, capturing biologically meaningful similarity beyond sequence identity. Furthermore, they suggest that CLAlign’s contrastive learning framework not only improves alignment accuracy but also yields a scoring function that better encodes structural and evolutionary correspondence, enabling accurate homology detection even in the absence of structural information.

### 2.5 CLAlign extends to structure alignment with high efficiency

CLAlign accommodates multiple input types through interchangeable encoders. To enable direct structure-based alignment, the sequence encoder can be replaced with the structure-informed encoder from ESM-IF [26], yielding CLAlign-ESMIF. CLAlign-ESMIF operates on 3D coordinates without modifying the overall architecture or the training objective, and it produces substitution matrices and alignments through the same downstream dynamic programming. This modularity enables sequence-based and structure-based alignment to be treated as instances of a single learning-and-inference pipeline.

CLAlign-ESMIF was evaluated on MALIDUP and MALISAM against representative structure-based methods, including FAST, TM-align, Dali, and GTAlign. Across both datasets, CLAlign-ESMIF achieves residue-level F1-scores comparable to or higher than those of TM-align and GTAlign, while maintaining competitive TM-scores (Table S2 and Fig. S2). These results suggest that CLAlign-ESMIF yields more biologically consistent residue correspondences even when global geometric similarity is slightly lower, offering an efficient and scalable alternative for large-scale structural comparison with minimal changes to the CLAlign framework.

### 2.6 Case study

To illustrate how contrastive learning improves alignment quality, a representative MALISAM pair, d1cs1a-d1q8ia1 (SCOPe IDs: d1cs1a and d1q8ia1), was examined. The manually curated reference alignment for this pair yields a TM-score of 0.576. In comparison, the sequence-based pLM methods pLM-BLAST, PLMAlign, and EBA achieve substantially lower TM-scores (0.269, 0.309, and 0.238, respectively), whereas CLAlign increases the TM-score to 0.493, approaching the reference alignment.

The improvement is more pronounced at the residue-correspondence level. The F1-scores of pLM-BLAST, PLMAlign and EBA are 0.151, 0.158, and 0.196, respectively, indicating that only a small fraction of the correct residue pairings are recovered by these methods. By contrast, CLAlign attains an F1-score of 0.569, representing a substantial increase in residue-level correctness.

Fig. 4 further visualizes structural alignment results and the substitution matrices produced for d1cs1a-d1q8ia1. CLAlign aligns the alpha-helices almost perfectly with the manual reference, whereas other sequence-based methods exhibit deviations. For beta-strands, other methods show small mis-alignments. In addition, pLM-BLAST, PLMAlign, and EBA show noticeable deviations in loop and terminal regions, whereas CLAlign more closely follows the manual reference alignment. These visual differences support the quantitative results and indicate that CLAlign provides more structure-aware and residue-consistent alignments. Matrices derived from pLM-BLAST, PLMAlign, and EBA exhibit diffuse similarity patterns without a coherent diagonal trajectory, making it difficult for dynamic programming to separate aligned from non-aligned positions. In contrast, CLAlign produces a sharply defined alignment path, increasing the margin between true correspondences and background similarity. The improved matrix structure, together with the increases in TM-score and F1-score, supports the interpretation that contrastive fine-tuning yields more structure-aware representations and thereby improves alignment accuracy.

**Figure 4.**
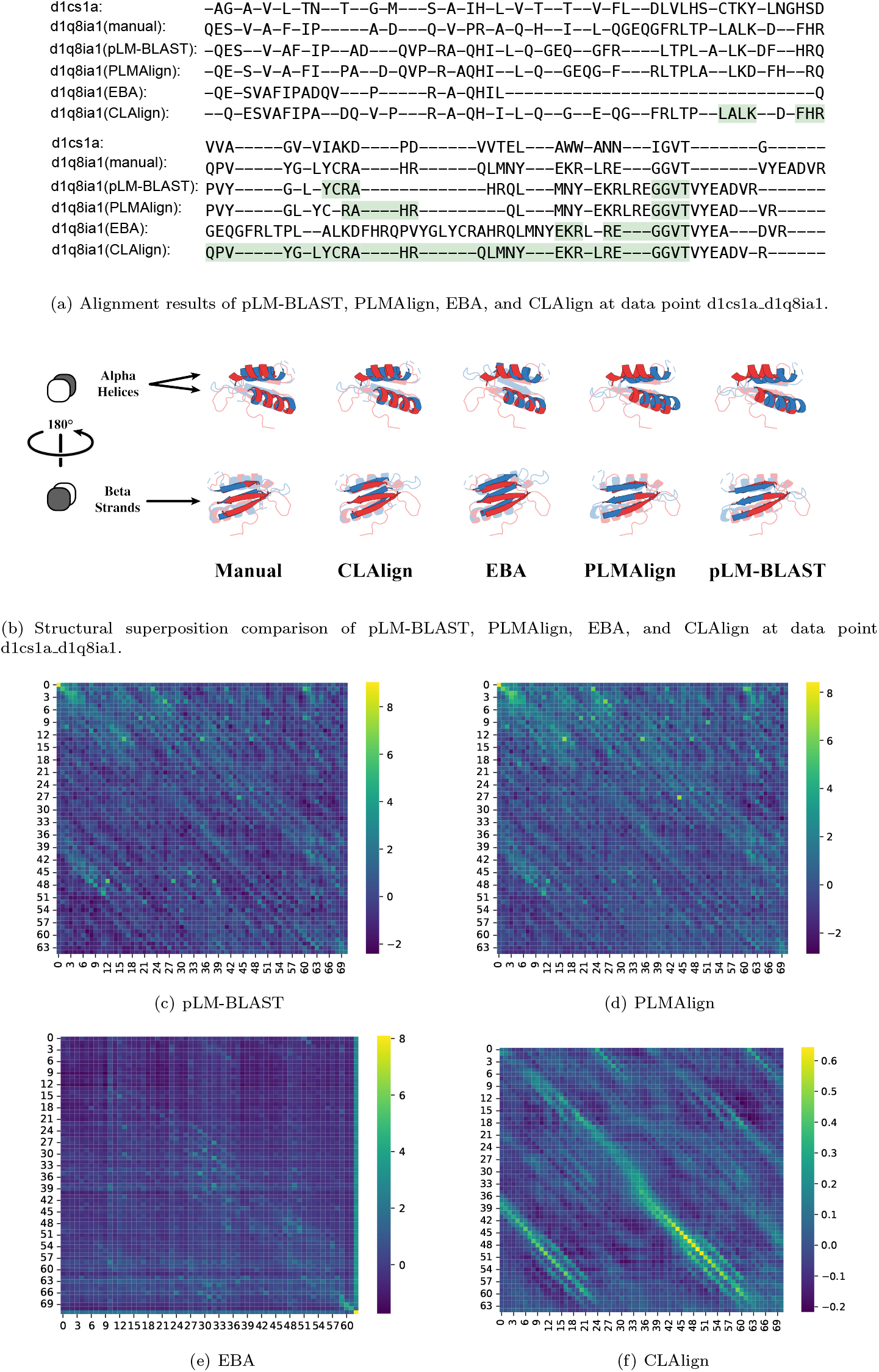
(a) Alignment results, (b) structural superposition comparison, and heatmap of substitution matrices of (c) pLM-BLAST, (d) PLMAlign, (e) EBA, and (f) CLAlign for data point d1cs1a d1q8ia1 (SCOPe IDs: d1cs1a and d1q8ia1) from MALISAM. Only CLAlign’s heatmap contains a clear alignment trajectory.

### 2.7 Running Time

CLAlign is trained on a relatively small dataset (10,000 samples), enabling rapid fine-tuning: training completes in 12 min on a system with 4 *×* 4090 NVIDIA GPUs. During inference, CLAlign follows the same overall workflow as embedding-based sequence aligners such as pLM-BLAST, PLMAlign, and EBA, resulting in comparable per-pair costs. Through efficient multi-core parallelization, however, CLAlign achieves substantially higher throughput in large-scale alignment scenarios.

Runtime was benchmarked on the SCOPe40-test dataset using a cluster equipped with dual Intel(R) Xeon(R) Platinum 8352V CPUs @ 2.10 GHz (72 cores, 144 threads) and 512 GB of RAM. SCOPe40-test contains 2,207 proteins, corresponding to 4,870,849 all-versus-all pairwise alignments. After database preprocessing, CLAlign completes the full set of alignments in 11-13 min. In comparison, GTAlign requires 50 min under default settings. Overall, CLAlign is more than fourfold faster than GTAlign, even though GTAlign is the most parallelized program among the structure-based methods. TM-align and Dali run slower than GTAlign, reflecting higher algorithmic complexity and the lack of native parallelization. Together, these results highlight that CLAlign combines high alignment accuracy with practical efficiency for large-scale analysis in both sequence-only and structure-informed settings.

## 3 Discussion

CLAlign highlights the utility of structure-aware residue representations for protein alignment. By contrastively fine-tuning protein language models, CLAlign consistently surpasses traditional sequence-based methods and recent deep learning sequence aligners, including pLM-BLAST, PLMAlign, EBA, and DeepBLAST. Performance was evaluated on two complementary benchmarks: MALIDUP, which emphasizes homologous domain pairs generated by internal duplication, and MALISAM, which contains analogous motifs without common ancestry. The strongest gains are observed on MALISAM, where accurate mapping depends primarily on structural similarity rather than sequence identity. Beyond accuracy, the results suggest that contrastive supervision can convert general-purpose pLM embeddings into alignment-specific representations that are more amenable to classical dynamic programming. Unlike prior methods such as DeepBLAST and DEDAL, which directly optimize alignment via differentiable dynamic programming, CLAlign instead focuses on learning better residue representations. By contrastively pulling matched residues together and pushing mismatched ones apart, it enables standard dynamic programming to recover more accurate alignments.

In structure-based alignment, CLAlign achieves higher F1-scores than TM-align, indicating more accurate residue-level correspondences. Although TM-align may achieve higher TM-scores, the F1-score is more biologically meaningful and serves as our primary metric. These results demonstrate that CLAlign can effectively leverage structural information to produce high-quality alignments.

A notable result is that CLAlign achieves alignment quality comparable to TM-align using sequence input alone. This suggests that contrastively trained protein language models encode structural information implicitly, enabling accurate alignment without explicitly providing 3D coordinates of structural data. Importantly, this sequence-only mode also offers practical advantages. CLAlign has lower computational complexity and substantially faster runtime than TM-align, which makes it well-suited for large-scale applications such as proteome-wide comparisons and high-throughput homology searches.

A gap nevertheless remains between sequence-only and structure-informed alignment. Closing this gap is an important direction for future work and may benefit from richer contrastive objectives, larger and more diverse supervision sets, or the integration of predicted structural features such as inter-residue geometry, confidence-weighted restraints, or contact probabilities. Such extensions could further strengthen the ability of sequence-only representations to recover reliable structural correspondence while maintaining the computational efficiency required for proteome-scale applications.

## 4 Method

### 4.1 Protein language model

Pretrained language models based on the Transformer architecture [27] have not only achieved tremendous success in natural language processing tasks [28] but have also made significant strides in the field of bioinformatics [29, 11, 13–15, 26]. The core idea of the Transformer structure is to utilize the self-attention mechanism to weigh the importance of different tokens relative to each other and capture complex relationships between them. Currently, pretrained language models based on protein sequences, such as TAPE [29], ESM [11, 13], and ProtT5 [14], have improved the performance of various protein downstream tasks, including structure prediction, homology detection, and functional annotation.

In CLAlign, we also use a protein pretrained language model (pLM) as the encoder for proteins. Suppose that we have a *N* -length protein sequence or structure *X* = *{x*_1_, *x*_2_, …, *x*_*N*_ *}* and a *M* -length protein sequence or structure *Y* = *{y*_1_, *y*_2_, …, *y*_*M*_ *}*, the matched *T* residues of *X* and *Y* are 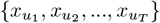 and 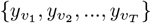, respectively. We use *f*_PLM_ to gain embeddings of *X* and *Y* as follows:

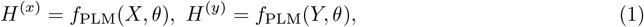

where *θ* is the trainable parameters of *f*_PLM_, *d* is the dimension of embeddings, *H*^(*x*)^ *∈ℝ*^*N ×d*^ and *H*^(*y*)^ *∈ℝ*^*M ×d*^ are embeddings of *X* and *Y*, respectively.

For sequence alignment, we employ ProtT5 [14] or ProstT5 [15] as the pLM encoder. Both ProtT5 and ProstT5 are based on the T5 architecture [30]. ProtT5 is trained solely on protein sequences using a conventional masked language modeling (MLM) objective. In contrast, ProstT5 is trained with a sequence–structure translation objective, enabling it to generate sequence representations enriched with protein structural semantics. For structure alignment, we adopt the encoder component of ESM-IF [26] as the pLM encoder. This encoder integrates geometric vector perceptron (GVP) layers with standard transformer layers, directly modeling the 3D coordinates of protein atoms to produce deep semantic representations of amino acids that are rich in structural information.

### 4.2 Contrastive Learning

Then we define contrastive loss function *ℓ* for the *i*-th matched residues as follows:

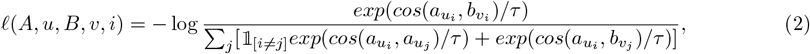

where *a*_*i*_, *b*_*i*_ are the *i*-th row vector of the embedding matrix *A* and *B*, respectively, and *τ* is the temperature parameter. The final contrastive loss function *ℒ* is calculated as follows:

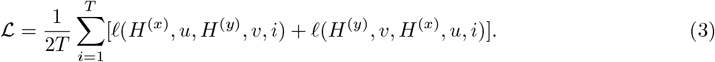

The extension of the contrastive loss function to batch sizes greater than one is equivalent to computing the loss on a single concatenated pair, thereby preserving the original training objective.

### 4.3 Alignment

After contrastive learning, we obtain the substitution matrix from the structure-aware embeddings and use NW to generate the alignment results, where the process is similar to pLM-BLAST and PLMAlign. For details, each element *s*_*i,j*_ of substitution matrix *S ∈ ℝ*^*N ×M*^ for protein *X* and *Y* is defined as

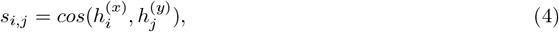

where 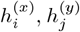 are the *i*-th row vector of *H*^(*x*)^ and the *j*-th row vector of *H*^(*y*)^, respectively, corresponding to the embeddings of the *i*-th amino acid of protein *X* and the *j*-th amino acid of protein *Y* . Then, we use a dynamic programming algorithm based on the substitution matrix *S* to find the alignment with the highest score. The dynamic programming recurrence relation is as follows:

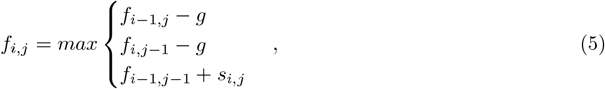

where *g* is the gap penalty, *f*_*i,j*_ represents the maximum score for matching the first *i* amino acids of protein *X* with the first *j* amino acids of protein *Y*, and *f*_*N,M*_ is the score of the alignment between protein *X* and *Y* . We obtain the alignment by backtracking through the transition process of *f*_*N,M*_ . All CLAlign results reported in this study use the NW formulation described above. Because SW uses the same substitution matrix and differs only in the standard local-alignment dynamic-programming boundary and traceback conditions, it is not described separately.

### 4.4 Data collection

We conducted structural alignment benchmarks using the two common databases, MALIDUP [23] and MALISAM [24]. MALIDUP consists of 241 manually curated pairwise structural alignments of duplicated domain pairs, while MALISAM comprises 130 pairwise structural alignments of structurally analogous motifs in proteins.

We used the SCOPe40-test dataset provided by PLMAlign [17] as the test set for remote homology detection. The SCOPe (Structural Classification of Proteins-extended) dataset [31, 32] organizes proteins into a hierarchical classification system based on their structural similarities and evolutionary relationships. It categorizes proteins into families, superfamilies, and folds, reflecting their homologous relationships. This structured approach allows researchers to explore the connections between proteins and understand their functional and structural diversity more effectively. The SCOPe40-test dataset is constructed from SCOPe40 version 2.01, which contains 2207 proteins.

To construct the training set for CLAlign, we retain proteins from DeepBLAST’s training set that simultaneously met the following conditions: (1) a length of no more than 510 amino acids; (2)a TM-score computed with TM-align between 0.3 and 0.9; (3) an e-value greater than 10.0 with the proteins in MALIDUP, MALISAM, and SCOPe40-test according to phmmer^1^. This ensures that the resulting training set is minimally similar to the test sets. Ultimately, we obtained around 300,000 pairwise structural alignments from the DeepBLAST’s training set. To accelerate training, we randomly sampled 10,000 pairs from these 300,000 training pairs as the final training dataset. During the sampling process, we prioritized selecting pairs from different proteins, so that the final 10,000 pairs included as many distinct proteins as possible. We constructed the validation set in the same manner as the training set from DeepBLAST’s validation data, except that TM-scores of protein pairs were restricted to the range 0.6–0.9 to avoid extremely dissimilar pairs. The validation set was then subsampled to 500 pairs.

### 4.5 Baselines

We compare CLAlign with two traditional methods: Needleman-Wunsch(NW) and Smith-Waterman(SW), and five PLM-based methods: three unsupervised methods pLM-BLAST [16], PLMAlign [17] and EBA [18], and two supervised methods DeepBLAST [19] and DEDAL [20]. We also compare CLAlign with five structure-based methods: Mammoth-local [33], FAST [3], TM-align [4], Dali [5], and GTAlign [6].

### 4.6 Evaluation metrics

We evaluate alignment quality using residue-level accuracy, global structural similarity, and secondary-structure consistency (HEC). For metrics requiring length normalization, we use the average sequence length

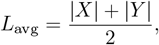

where |*X*| and |*Y* | denote the lengths of the two proteins.

#### Residue-level accuracy

For a given protein pair, let *S* = *{*(*u*_1_, *v*_1_), …, (*u*_*m*_, *v*_*m*_)*}* denote the predicted aligned residue pairs, and 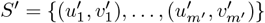 denote the reference alignment. Precision, Recall, and F1-score for this pair are defined as:

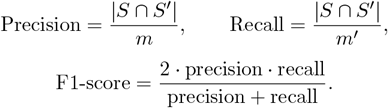

For each benchmark, the reported precision, recall, and F1-score are obtained by averaging these per-pair quantities over all protein pairs.

#### Global structural similarity

To measure global geometric consistency, we compute TM-score using *L*_avg_:

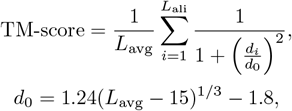

where *L*_ali_ is the number of aligned residue pairs and *d*_*i*_ is the C^*α*^ distance between the *i*-th aligned residues.

#### Secondary-structure consistency (HEC). HEC-ACC

Let 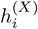 and 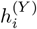 be the H/E/C states of aligned residues. We normalize by *L*_avg_:

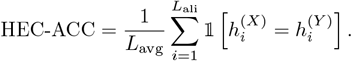

#### HEC-SOV (segment overlap)

To quantify the segment-level consistency of secondary structure under the predicted alignment, we compute a simplified SOV score defined as follows.

#### Notation

For each protein *X* = (*x*_1_, …, *x*_|*X*|_), let 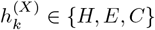 denote the secondary-structure state of residue *x*_*k*_. A secondary-structure segment of type *t ∈ {H, E}* is a maximal interval

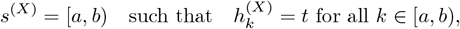

and satisfying the minimal length constraint

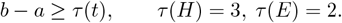

This yields the segment sets *S* ^(*X*)^ and *S*^(*Y*)^ for the two proteins.

Let *m*(*k*) denote the alignment mapping:

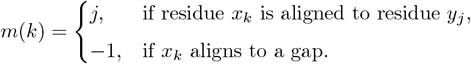

**Overlap count**. For a pair of segments *s*^(*X*)^ = [*a, b*) *∈ S*^(*X*)^ and *t*^(*Y*)^ = [*c, d*) *∈ S*^(*Y*)^ with the same secondary-structure type, we define the overlap count

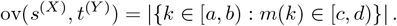

This counts how many residues in *s*^(*X*)^ are aligned into *t*^(*Y*)^.

#### Best-overlap ratios

For each segment 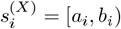 in *X*, we define

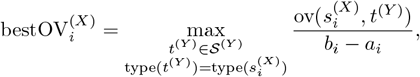

i.e. the maximum fraction of residues in 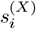 that are aligned into any segment of the same type in *Y* . Symmetrically, for each 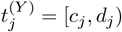 we define

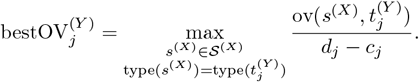

#### Final score

The final HEC-SOV score is the symmetric average

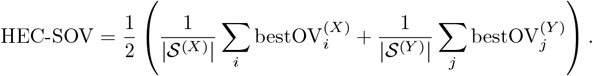

## 4.7 Data availability

MALIDUP is available at http://prodata.swmed.edu/malidup. MALISAM is available at http://prodata.swmed.edu/malisam. SCOPe40 is available at https://scop.berkeley.edu/.

## 4.8 Code availability

The source code is available at https://github.com/yourh/CLAlign.

## Footnotes

1 https://github.com/EddyRivasLab/hmmer

## Notes

### Competing Interest Statement

The authors have declared no competing interest.

### Summary of Updates

This version has been revised to provide a more detailed description of the proposed method, add additional experiments, and expand the manuscript content. New experimental results and analyses have been included to further support the main findings. The presentation, discussion, and overall organization of the manuscript have also been improved.

