## supplementary materials for "Effective structure-aware protein alignment via residue-level contrastive learning"

#### 1 Experimental Settings

We implement CLAlign in Python with PyTorch [1] and Transformers [2]. For a fair comparison, we employ ProtT5 [3] (prot_t5_xl_uniref50) as the pLM encoder of CLAlign, which is the same as DeepBLAST, pLM-BLAST and PLMAlign. We also evaluated two variants: CLAlign-ProstT5, which uses ProstT5 [4] for enhanced performance, and CLAlign-ESMIF, which uses the encoder of ESM-IF [5] for structure alignment. We fine-tuned the pLM using LoRA [6] with rank 8, optimized with Adam [7] at a learning rate of $2e-5$. Training was performed for 3 epochs with a batch size of 8. The temperature parameter $\tau$ we used in contrastive learning is 0.1. We set the gap penalty $g$ to 0.0 directly, as the pLM-based substitution matrix already captures contextual similarity, making additional gap penalties unnecessary.

All baselines were run with their default settings, including gap penalties. For alignment quality assessment, global alignments from all baseline methods were used, except for Smith–Waterman (SW) and SW-based DEDAL, which only support local alignments.

#### 2 The Impact of Gap Penalty

As shown in Fig. S1, the F1-scores of all three datasets, validation set, MALIDUP, and MALISAM, decrease as the gap penalty increases, which supports setting the gap penalty to 0.0 when using the pLM-based substitution matrix.

#### 3 Performance Evaluation of CLAlign Across Multiple Runs

Table S1 summarizes the precision, recall, and F1-score of CLAlign across five independent runs. The results show that the performance is consistent, with very small standard deviations across runs, indicating that the method is stable.

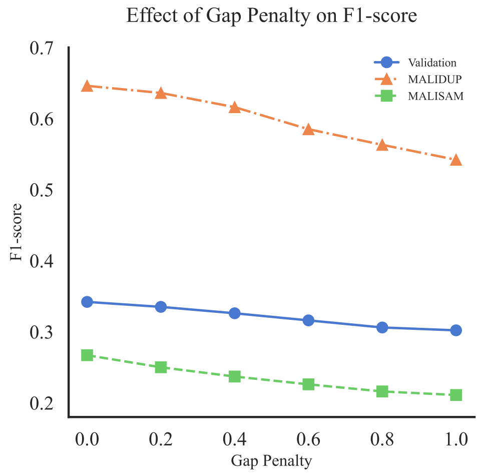


Figure S1: Effect of gap penalty on F1-score across Validation set, MALIDUP and MALISAM. All datasets show a decreasing trend as the gap penalty increases.

Table S1: Performance of CLAlign over five independent runs on three datasets: Validation, MALIDUP, and MALISAM. The first row (avg) reports the mean values of precision, recall, and F1-score, while the second row (std) shows the corresponding standard deviations, indicating the stability of the model across runs.

| $i$th | Validation | | | MALIDUP | | | MALISAM | | |
|---|---|---|---|---|---|---|---|---|---|
| | Precision | Recall | F1-score | Precision | Recall | F1-score | Precision | Recall | F1-score |
| 1 | 0.336 | 0.349 | 0.342 | 0.642 | 0.650 | 0.646 | 0.261 | 0.274 | 0.267 |
| 2 | 0.332 | 0.344 | 0.338 | 0.640 | 0.648 | 0.644 | 0.260 | 0.273 | 0.266 |
| 3 | 0.334 | 0.347 | 0.340 | 0.637 | 0.643 | 0.640 | 0.259 | 0.271 | 0.265 |
| 4 | 0.332 | 0.345 | 0.338 | 0.637 | 0.645 | 0.641 | 0.259 | 0.271 | 0.264 |
| 5 | 0.332 | 0.345 | 0.338 | 0.639 | 0.646 | 0.642 | 0.257 | 0.270 | 0.263 |
| avg | 0.333 | 0.346 | 0.339 | 0.639 | 0.646 | 0.643 | 0.259 | 0.272 | 0.265 |
| std | 0.004 | 0.002 | 0.002 | 0.002 | 0.002 | 0.002 | 0.002 | 0.002 | 0.002 |

Table S2: Performance comparison of protein structure alignment on MALIDUP and MALISAM

| Methods | MALIDUP | | | MALISAM | | |
|---|---|---|---|---|---|---|
| | Precision | Recall | F1-score | Precision | Recall | F1-score |
| Mammoth-local | 0.520 | 0.522 | 0.520 | 0.220 | 0.231 | 0.225 |
| FAST | **0.875** | 0.714 | 0.781 | **0.654** | 0.493 | 0.559 |
| TM-align | 0.755 | 0.709 | 0.730 | 0.478 | 0.441 | 0.458 |
| Dali | 0.800 | **0.791** | **0.795** | 0.630 | **0.630** | **0.630** |
| GTAlign | 0.749 | 0.710 | 0.727 | 0.505 | 0.475 | 0.489 |
| CLAlign-ESMIF | 0.722 | 0.722 | 0.722 | 0.521 | 0.531 | 0.526 |

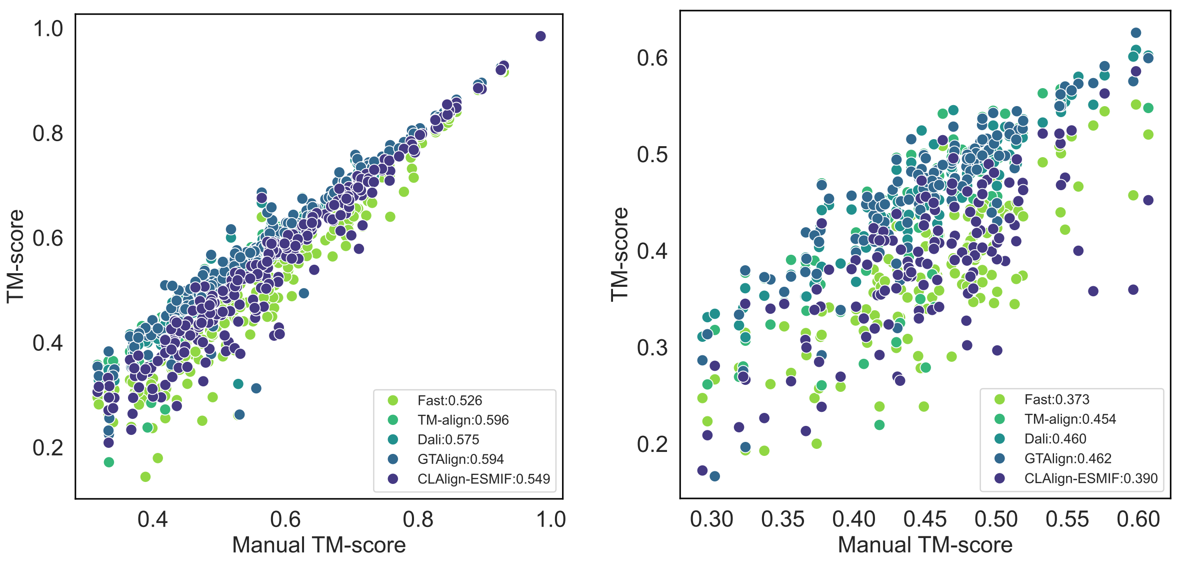


(a) Comparison of structure alignment on MALIDUP

(b) Comparison of structure alignment on MALISAM

Figure S2: Comparison of global structural similarity (TM-score) among different structure methods on (a) MALIDUP and (b) MALISAM. The x-coordinate of each point represents the TM-score of the manual alignment results for that data point, while the y-coordinate represents the TM-score of the alignment results obtained by the compared methods. For each method in the legend, the number after the colon represents its average TM-score.

### 4 Comparison of CLAlign with structure-based methods

Table S2 summarizes the precision, recall, and F1-score of CLAlign-ESMIF and representative structure alignment methods on MALIDUP and MALISAM. For Dali on MALIDUP, seven failed alignments were assigned zero scores when computing the aggregate metrics reported in Table S2 and Fig. S2. CLAlign-ESMIF demonstrated competitive performance relative to established structure alignment methods. In terms of residue-level F1-score, Dali achieves the best results among structure alignment methods, with FAST performing slightly lower. CLAlign-ESMIF is lower than Dali and FAST but remains comparable to TM-align and GTAlign on MALIDUP, while outperforming TM-align and GTAlign on MALISAM by 14.8% and 7.6%, respectively. These results position CLAlign-ESMIF alongside the strongest structure-based methods, surpass several widely used tools, and underscore its effectiveness in challenging cases of structural analogy without sequence homology.

Fig. S2 presents TM-score comparisons among five protein structure methods: FAST, TM-align, Dali, GTAlign, and CLAlign-ESMIF. Dali, TM-align, and GTAlign achieved relatively high TM-scores, while CLAlign-ESMIF scored below them and FAST scored slightly below CLAlign-ESMIF. CLAlign-ESMIF showed decreases of approximately 7.9% on MALIDUP and 14.1% on MALISAM relative to TM-align. However, CLAlign-ESMIF remains comparable to TM-align in F1-score on MALIDUP and achieves a substantially higher F1-score on MALISAM, indicating that it can produce biologically meaningful alignments despite lower TM-scores.
